# The consequences of variant calling decisions in secondary analyses of cancer genomics data

**DOI:** 10.1101/2020.01.29.924860

**Authors:** Carlos García-Prieto, Francisco Martínez-Jiménez, Alfonso Valencia, Eduard Porta-Pardo

## Abstract

The analysis of cancer genomes provides fundamental information about its aetiology, the processes driving cell transformation or potential treatments. The first crucial step in the analysis of any tumor genome is the identification of somatic genetic variants that cancer cells have acquired during their evolution. For that purpose, a wide range of somatic variant callers have been developed in recent years. While there have been some efforts to benchmark somatic variant calling tools and strategies, the extent to which variant calling decisions impact the results of downstream analyses of tumor genomes remains unknown.

Here we present a study to elucidate whether different variant callers (MuSE, MuTect2, SomaticSniper, VarScan2) and strategies to combine them (Consensus and Union) lead to different results in these three important downstream analyses of cancer genomics data: identification of cancer driver genes, quantification of mutational signatures and detection of clinically actionable variants. To this end, we tested how the results of these three analyses varied depending on the somatic mutation caller in five different projects from The Cancer Genome Atlas (TCGA). Our results show that variant calling decisions have a significant impact on these downstream analyses, creating important differences in driver genes identification and mutational processes attribution among variant call sets, as well as in the detection of clinically actionable targets. More importantly, it seems that Consensus, a very widely used strategy by the research community, is not the optimal strategy, as it can lead to the loss of some cancer driver genes and actionable mutations. On the other hand, the Union seems to be a legit strategy for some downstream analyses with a robust performance overall.

## 1 Introduction

The widespread access to genomic data of tumor samples and cancer patients is transforming all aspects of this disease, from basic research to its clinical care^1^. For example, thanks to sequencing data, we are beginning to understand the aetiology of the mutational processes that affect cancer cells^2^. Furthermore, since cancer progression relies on the expansion of clonal cell populations, we are now able to track and reconstruct the phylogenetic tree of tumor evolution^3^. Similarly, the large cohorts of cancer patients that have been sequenced so far, have helped us identify germline and somatic mutations that predispose or drive carcinogenesis respectively^4,5^, laying the foundation of personalized cancer care.

The first crucial step in analyzing cancer genomic data is the identification of genetic variants, particularly those of somatic origin. In that sense, the research community has made great efforts to assess the performance of the many different somatic variant callers available^6–11^. However, so far, there has been no agreement on which variant caller nor which strategy to combine them is the most suitable. For instance, The Cancer Genome Atlas (TCGA) implemented different variant callers on multiple papers throughout its history^12–15^. This eventually led to the Multi-Center Mutation Calling in Multiple Cancers (MC3) project^16^ to address standardization and reproducibility issues at the end of TCGA. During MC3, many groups worked together to define a clear and unique strategy to combine the output of multiple variant calling tools. Another way to combine the output of different tools is to use machine learning approaches^17,18^. However, despite all these efforts, it is still unclear which variant calling tool, or combination of tools, should be used to analyze cancer genomics data.

While the biggest challenge in determining the optimal variant calling tool or strategy is the lack of gold standard sets of somatic variants, another likely important reason is that it is difficult to define what is actually best in cancer genomics. At the end of the day somatic variant calling is a means to an end. Sequencing data from tumors can be used for many different secondary analyses, from finding cancer driver genes and mutations to determining the presence of clinically actionable mutations or quantifying the effects of mutational signatures. Since none of the somatic variant callers nor strategies are perfect, it is possible that the answer to all these secondary analyses differs depending on which somatic variant calling tool or strategy is used.

While there have been benchmarking studies comparing how mutation callers find somatic mutations, to the best of our knowledge there has been no systematic study of the impact on variant calling tools in secondary analyses. In this paper we studied how decisions at the somatic variant calling stage of cancer genomics data, affect the results of three different secondary analyses: detection of cancer driver genes and mutations, quantification of mutational signatures and identification of clinically actionable variants.

## 2 Methods

### Variant calling datasets

To compare the effects of different mutation calling approaches in secondary analyses, we selected five different projects from TCGA: adrenocortical carcinoma (ACC)^19^, bladder urothelial carcinoma (BLCA)^20^, breast invasive carcinoma (BRCA)^13,21^, prostate adenocarcinoma (PRAD)^22^ and uterine corpus endometrial carcinoma (UCEC)^23^. We chose these five projects to explore the impact of the somatic variant calling strategy in downstream analyses of cohorts with different sizes, mutational signatures and mutational burdens. The Genomic Data Commons (GDC) portal (https://portal.gdc.cancer.gov) gives access to all the processed whole-exome sequencing (WXS) data for all the TCGA projects. In particular, the GDC created the DNA-Seq pipeline to process all TCGA samples in a uniform way^24^. This pipeline includes sample preprocessing, alignment to the human reference genome GRCh38.d1.vd1 followed by BAM cleaning and somatic variant calling with variant annotation and aggregation. Somatic variants were identified in WXS data by comparing allele frequencies in matched tumor-normal samples. The GDC used four different variant calling tools to identify somatic mutations: MuSE^25^, MuTect2^26^, SomaticSniper^27^ and VarScan2^28^. After analyzing the whole-exome sequencing data for each individual sample, the GDC pipeline includes an aggregation step that combines variants from all cases of a cancer cohort into a single TCGA project mutation annotation format (MAF) file.

For each of the five different cancer types, we downloaded the four different aggregated MAF files with all the somatic mutations for each variant caller. Additionally, we computed two extra mutation call sets per TCGA project. First, we created a Consensus file with those variants that were called by at least 3 out of the 4 aforementioned variant callers. Finally, we also created a Union file with every somatic variant called by any variant caller.

Regarding variant calling tools, we must point out that only MuTect2 and VarsCan2 were able to call small indels. Moreover, this also forced the Consensus call set to include only SNVs. Overall, indels represented less than 10% of variants called by each of these two variant callers. For this reason, the variant call sets analysed in the study were composed mainly of single nucleotide variants (SNVs).

### Detecting cancer driver genes

To identify cancer driver genes we used the intOGen pipeline (https://bitbucket.org/intogen/intogen-plus/src/master/, 03/20/2020)^29^. Specifically, we analyzed every somatic variant file (MuSE, MuTect2, SomaticSniper, VarScan2, Consensus and Union) of each of the five TCGA projects separately. IntOGen integrates the result of seven driver discovery methods: OncodriveFML^30^, OncodriveCLUSTL^31^, dNdScv^32^, CBaSE^33^, HotMAPS^34^, smRegions^35^ and MutPanning^36^. The driver discovery methods integrated in intOGen explore different signals of positive selection, such as clustering of mutations in protein structures or mutational functional bias, to pinpoint which driver genes deviate from the estimated neutral mutation rate using the set of input somatic mutations. The results of these tools are then combined by accounting each method credibility -the relative credibility for each method is based on the ability of the method to give precedence to well-known genes already collected in the Cancer Gene Census^37^ catalogue of driver genes- to produce a consensus ranking of genes using a TIER based classification. Finally, intOGen also provides a weighted combined p-value for each ranked gene.

For the purpose of our analysis, we only considered true driver genes those within TIER 1 and TIER 2 (q-value <0.05). We, therefore, discarded genes classified in TIER 3 and TIER 4.

### Benchmarking variant calling strategies with driver genes

We considered the curated set of known driver genes from intOGen (https://www.intogen.org/download, release date 2020/02/01) as our truth set to benchmark how the different mutation call sets can be used to detect cancer driver genes. This set encompasses both, newly detected and previously annotated cancer driver genes in the Cancer Gene Census (https://cancer.sanger.ac.uk/census) of Catalogue Of Somatic Mutations In Cancer (COSMIC)^38^.To further assess and compare our results, we also benchmarked against a second truth set comprised by the set of 299 driver genes published by the PanCancerAtlas -MC3- project^5^. We restricted our benchmarking analysis to known cancer driver genes in the cancer types we analyzed.

We used multiple metrics (**Table 1**) to assess the performance of the different variant calling strategies when detecting driver genes with intOGen in downstream analyses. We defined our True Positives (TP), False Positives (FP) and False Negatives (FN) as follows:

- TP: those driver genes detected by intOGen with a given variant call set that are within the truth set.
- FP: those driver genes detected by intOGen with a given variant call set that are outside the truth set.
- FN: those driver genes within the truth set not identified by intOGen with a given variant call set.

**Table 1.** Benchmarking metrics

| Metric | Description |
| --- | --- |
| $Precision = \frac{TP}{TP+FP}$ | Also known as Positive Predictive Value. It is the ratio of correctly detected driver genes among all driver genes detected by intOGen with a given somatic variant call set. |
| $Recall = \frac{TP}{TP+FN}$ | Also known as Sensitivity. It is the ratio of correctly detected driver genes by intOGen among all driver genes within the truth set. |
| $FDR = \frac{FP}{TP+FP}$ | It is the ratio of incorrectly detected driver genes among all driver genes detected by intOGen with a given somatic variant call set. |
| $F1 - Score = \frac{2*Precision*Recall}{Precision+Recall}$ | Harmonic average of precision and recall. The best value is 1 and the worst is 0. |

FDR: False Discovery Rate.

### Mutational signature analysis

We used deconstructSigs^39^ 1.8.0 R package to quantify the presence of different mutational signatures in the different mutation call sets. In brief, deconstructSigs accounts for the trinucleotide context of each mutation to classify the six different base substitutions (C>A, C>G, C>T, T>A, T>C, T>G) into 96 possible mutation types^2^. The signature matrix with the number of times a mutation was found within each trinucleotide context was compared against COSMIC Single Base Substitution (SBS) signatures (available at https://cancer.sanger.ac.uk/cosmic/signatures/SBS)^40^.

Finally, deconstructSigs uses an iterative approach to assign different weights to each signature and estimate their contribution to the mutational profile of the tumor sample. We filtered out those samples with less than 50 mutations. Since we analyzed WXS samples, the signature matrix was normalized to reflect the absolute frequency of each trinucleotide context as it would have taken place in the whole genome. This way we adjusted for differences in trinucleotide abundances between exome and whole genome in order to compare our signatures to the ones extracted from whole genomes (available in synapse.org, ID syn12009743).

To assess the different contribution of mutational signatures to a sample mutational profile we performed Kruskal-Wallis test comparisons across all variant call sets for a given cancer. Additionally, we also performed pairwise comparisons (Wilcoxon test) between one variant call set versus all the others for each signature and cancer.

### Clinically actionable variants analysis

We used the Cancer Genome Interpreter^41^ (CGI) (https://www.cancergenomeinterpreter.org, 2019/07/17) to detect alterations that might be therapeutically actionable. To detect alterations as biomarkers of drug response, CGI uses an in-house database of genomic events that influence tumor drug response (sensitivity, toxicity and resistance) classified with different levels of clinical evidence (FDA guidelines, case report, early trials, pre-clinical). Since CGI only supports hg19 mapped alterations, we transformed our MAF files to a BED-like format (keeping only “chr”, “start”, “end”, “ref allele”, “alt allele” and “sample ID” columns) and performed a liftover using CrossMap^42^ version 0.3.4 (99.99% of variants were successfully remapped).

### Purity dataset

We used purity/ploidy ABSOLUTE annotations^43^ for all TCGA samples available at https://gdc.cancer.gov/about-data/publications/pancanatlas.

### Clinical dataset

We retrieved tumor stage information from the TCGA-Clinical Data Resource^44^ file available at https://gdc.cancer.gov/about-data/publications/pancanatlas.

## 3 Results

### 3.1 Effects of variant calling in the detection of cancer driver genes

One of the most widespread uses of somatic mutation data from cohorts of cancer patients is the identification of cancer driver genes. The tools to detect cancer driver genes can be sensitive to which somatic mutations are included in the final analysis, as they can bias some aspects of the randomization in which most cancer driver detection tools rely to make their analysis^30–36^.

To assess to what extent decisions about calling somatic variants can affect the detection of cancer driver genes, we used intOGen^29^ to find cancer driver genes in thirty different mutation call sets for five different cancer types from TCGA. The six mutation call sets of each cancer type are distributed as follows: one mutation set with all the calls from one of the variant calling tools (MuSe^25^, MuTect2^26^, SomaticSniper^27^ and VarScan2^28^), another mutation set -Consensus- with the consensus of the four variant callers (i.e., all those mutations found by, at least, three of the four callers) and a final mutation set with all the mutations found by any mutation caller -Union- **(Figure 1)**.

**Figure 1.**
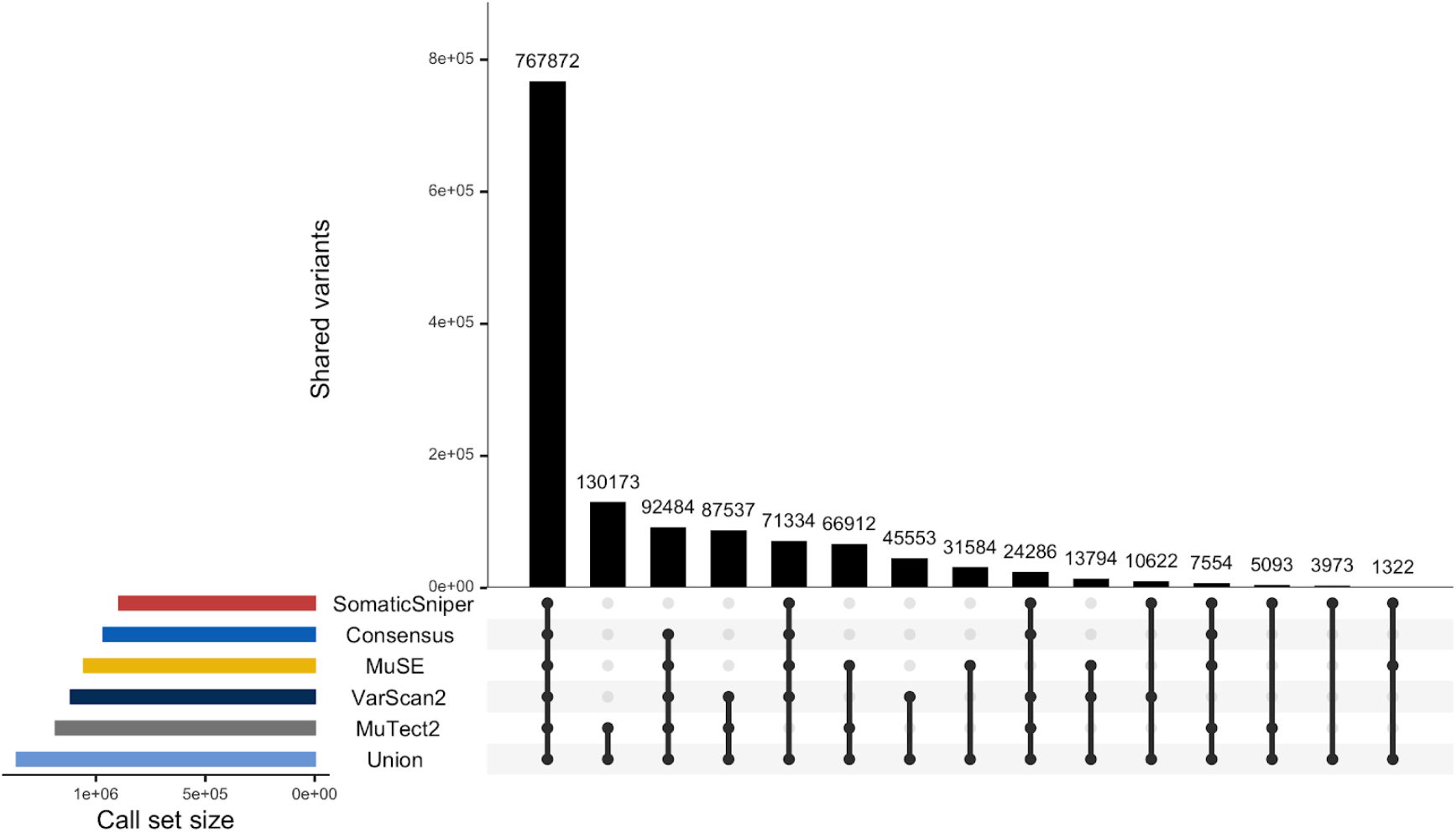
Intersection of mutation calls across variant calling strategies for the five cancer types together. This UpSetR plot shows the number of variants uniquely called by one tool (single point) and variants called by different tools (linked points). Top bar-plot indicates intersection size. Bottom left plot indicates call set size. See also Figure S2.

We chose five cancer types -adrenocortical carcinoma (ACC)^19^, bladder urothelial carcinoma (BLCA)^20^, breast invasive carcinoma (BRCA)^13,21^, prostate adenocarcinoma (PRAD)^22^ and uterine corpus endometrial carcinoma (UCEC)^23^= so that they spanned a variety of mutational processes, ranges of purity, mutation rates and cohort sizes within TCGA (**Figure S1**). For example, ACC is one of the smallest cohorts within TCGA (n = 92), as well amongst those with the highest tumor purity (average purity 80%)^45^. On the other hand, BRCA is the largest cohort in TCGA (n = 986). Another factor that can alter the efficiency of tools to detect cancer driver genes is the mutation rate of the cohort, hence why we included UCEC, which is amongst the cancer types with highest mutation rates^5^. Finally, BLCA and PRAD are amongst the cohorts that are closest to the TCGA average in all these aspects, making them good representatives of the average tumor sample.

In terms of absolute number of somatic mutations, the mutation call sets from SomaticSniper and MuTect2 were, respectively, the smallest and largest (not accounting for the Union) across all cancers with the exception of the ACC cohort (**Figure S2**). One possible factor affecting the number of somatic mutations identified by each caller is the sample purity. In fact, we found a positive correlation between the purity of a sample and the percent of all possible somatic variants (i.e.,Union call set) called by each variant caller, except for MuTect2, where the correlation was negative in all cancers (**Figure S3**). This shows that most of the mutations called in low purity samples were identified by MuTect2, suggesting that this tool has high sensitivity to identify somatic variants in low purity tumor samples^10,46^.

We next explored the relationship between sample purity and tumor stage. We found that only BLCA showed a correlation between purity and tumor stage: more advanced bladder tumors tend to be less pure (**Figure S4**). To further assess the role of tumor stage in the performance of somatic variant callers, we analysed the range of variant allele frequencies (VAFs) spanned by each call set (**Figure S5**). We observed that SomaticSniper call set spanned a median VAF range of 0.4-0.5 after adjusting for purity and ploidy^43^, whereas MuTect2 and Union call sets spanned lower VAF ranges, thus calling more subclonal mutations. This may suggest SomaticSniper as a reliable caller for early stage tumors, where we do not expect to find many subclonal populations. On the other hand, MuTect2 may be more suitable for late stage tumors.

Having assessed the influence of various tumor properties in the number of mutations called by each tool, we next wondered what the effect would be when trying to detect cancer driver genes. To that end, we used intOGen to detect cancer driver genes in the 30 somatic mutation call sets (**Figure 2, Table S1**).

**Fig. 2.**
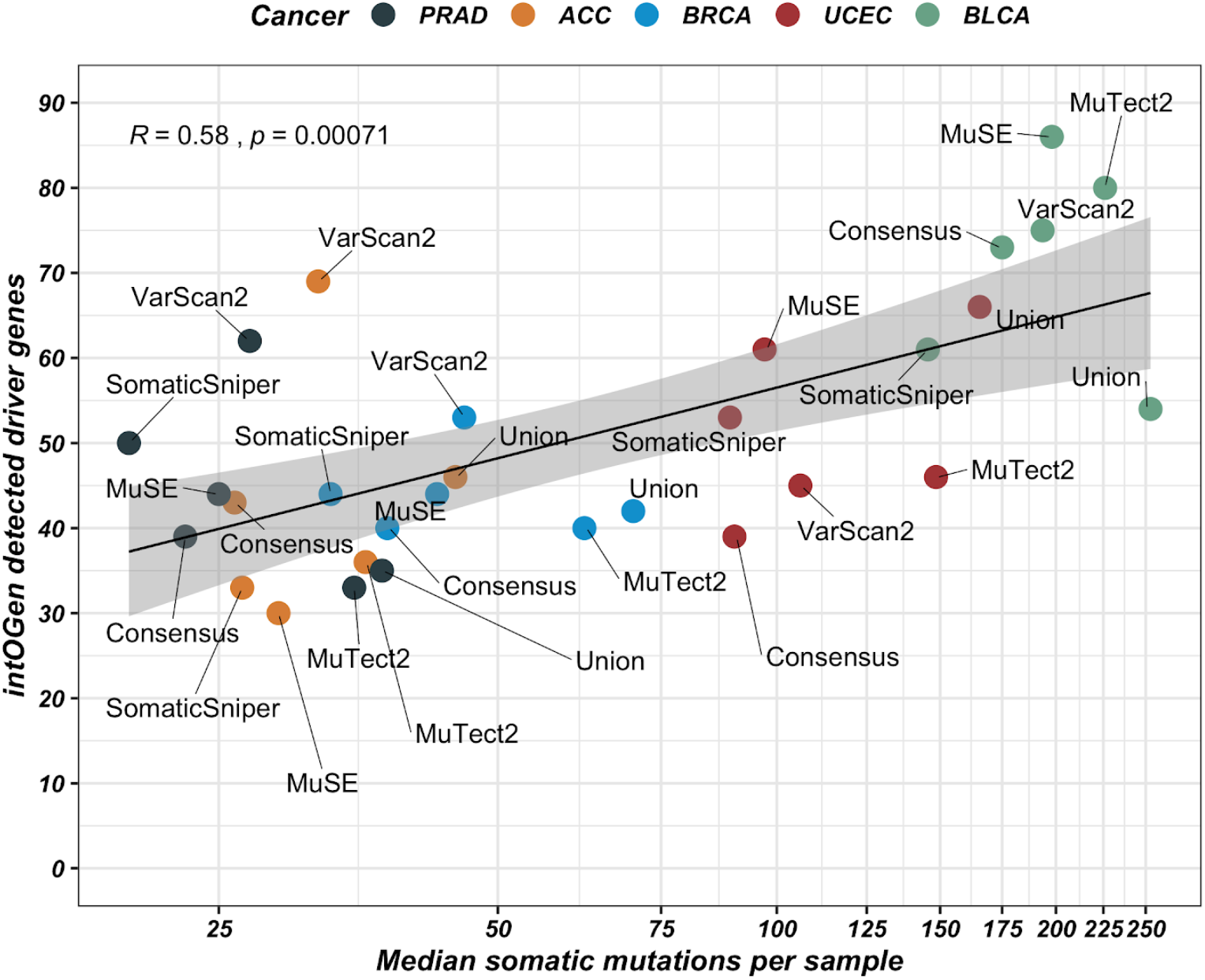
Correlation between cancer driver genes and median mutation burden. The number of cancer driver genes detected by intOGen with different call sets in each cancer type positively correlates with the median mutation burden. Shaded area indicates 95% bootstrapped confidence interval.

Overall, we found that there are wide differences in the number of detected cancer driver genes in each cohort depending on which somatic variant calling tool or strategy we used. For example in the case of prostate cancer, the set of mutations from MuTect2 leads to the detection of 33 cancer driver genes, whereas the set from VarScan2 leads to 62 driver genes. Similarly, in the case of bladder cancer, the Union leads to the detection of 54 cancer driver genes, whereas the set of mutations from MuSE leads to 86 driver genes. Interestingly, the number of cancer driver genes detected in each mutation call set has a positive correlation with the median mutation burden (spearman R = 0.58, p.value = 0.00071), as already described in the final driver analysis of TCGA^5^.

Next, we benchmarked our results against a gold standard set of known cancer driver genes from intOGen. We also considered the set of cancer driver genes published by the PanCancerAtlas -MC3- project^5^ as a second truth set to further assess our results. In both cases, we restricted our gold standard sets to only those genes annotated as cancer driver genes in the five tumor types we analysed. Thus, we considered five different and specific cancer type truth sets to classify the detected cancer driver genes as true positives, false positives and false negatives.

The aggregated benchmarking results of the five cancer types altogether showed, to our surprise, that the Union variant calling strategy is the best one when detecting cancer driver genes with intOGen (**Figure 3**). The Union call set is the top performer when benchmarking against both intOGen and MC3 truth sets, scoring 0.58 and 0.61 F1-Scores respectively. MuTect2 scores second best in both cases with very similar results (0.55 and 0.6 F1-Scores). The ranking of the different variant calling strategies performance is consistent when benchmarking against both truth sets.

**Fig. 3.**
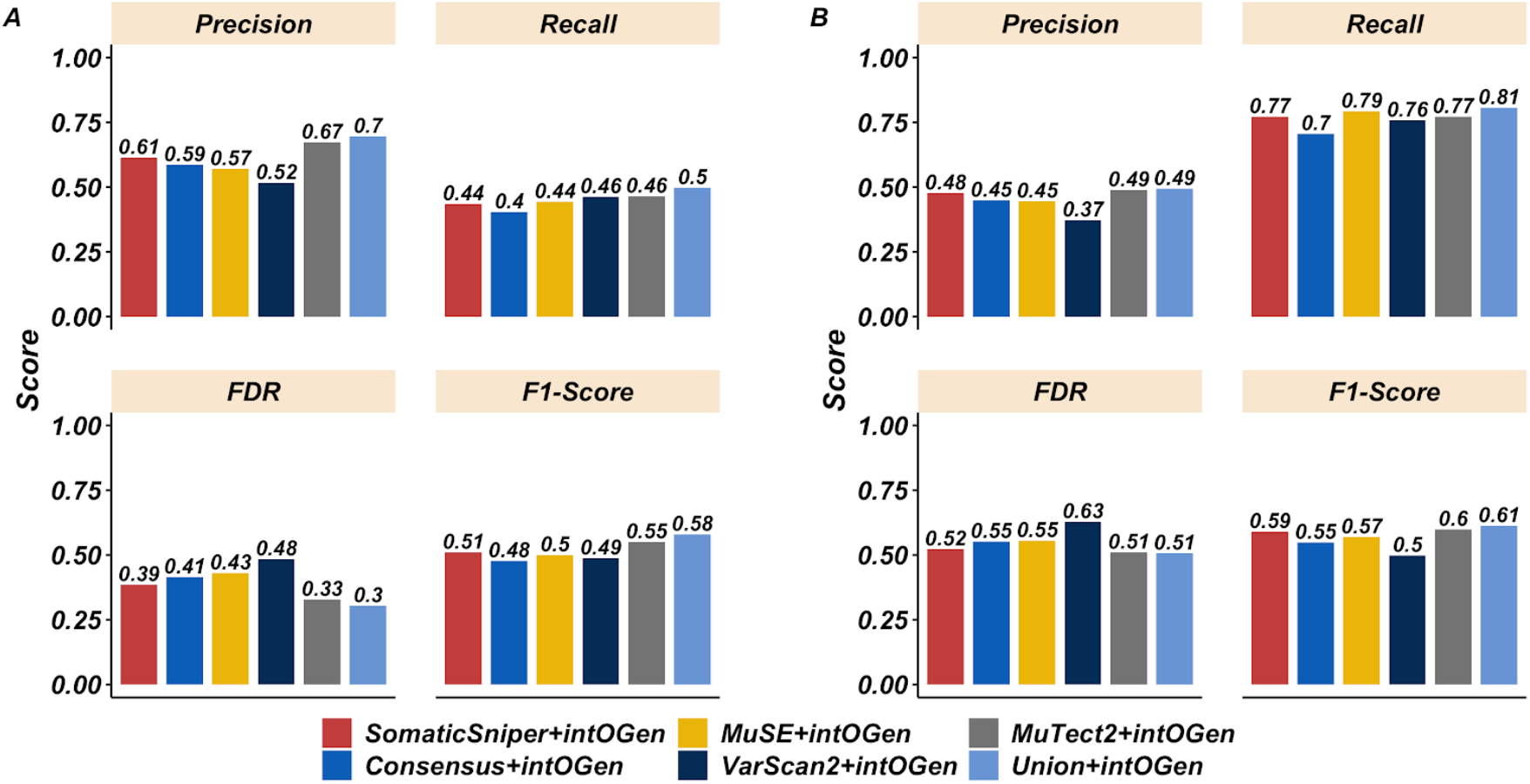
Performance metrics of the different variant calling strategies when detecting cancer driver genes with intOGen. The aggregated results for the five cancer types together are shown. (A) Performance metrics when benchmarking against int0Gen truth set of known cancer driver genes. (B) Performance metrics when benchmarking against PanCancerAtlas project -MC3- truth set of known cancer driver genes. FDR: False Discovery Rate. See also Figure S6.

One of the most widespread approaches to integrate the mutation call sets from different tools consists in generating a set with the consensus mutations. However, our results suggest that the resulting somatic mutation set is actually amongst the worst performers, alongside with that from VarScan2, to detect cancer driver genes. For example, the Consensus and Union F1-scores, 0.48 and 0.58 respectively, show that the latter has a better performance. Interestingly, the Consensus scores the lowest recall among all variant calling strategies. Furthermore, VarScan2 scores the highest false discovery rate among all variant calling strategies. On the other hand, SomaticSniper and MuSE scores rank in the middle of the different aforementioned performances.

The global performance of the different variant calling strategies is consistent with their benchmarking results at the individual cancer type level (**Figure S6**). The Union and MuTect2 calling methods continue to outperform the others, whereas the Consensus and VarScan2 results still are the worst ones. However, an exception to this trend is found in BRCA where the Consensus and Union performances are the same.

### 3.2 Somatic mutations in cancer driver genes

Even if one can identify a gene as a driver in a cohort using a variant call set, it is possible that some individual mutations in the gene in some samples are missing. This could have important implications for patients, as the presence or absence of mutations in cancer driver genes can determine whether patients will receive certain treatments or not.

To evaluate the impact of decisions about somatic variant calling when finding mutations in cancer driver genes, we calculated the ratio of patients with differing mutation status depending on the mutation set used. To this end, we accounted for the total number of samples in the Union call set with a mutation in a given driver gene to compute the percentage of those Union samples that were missing in each variant call set (**Figure 4**).

**Fig. 4.**
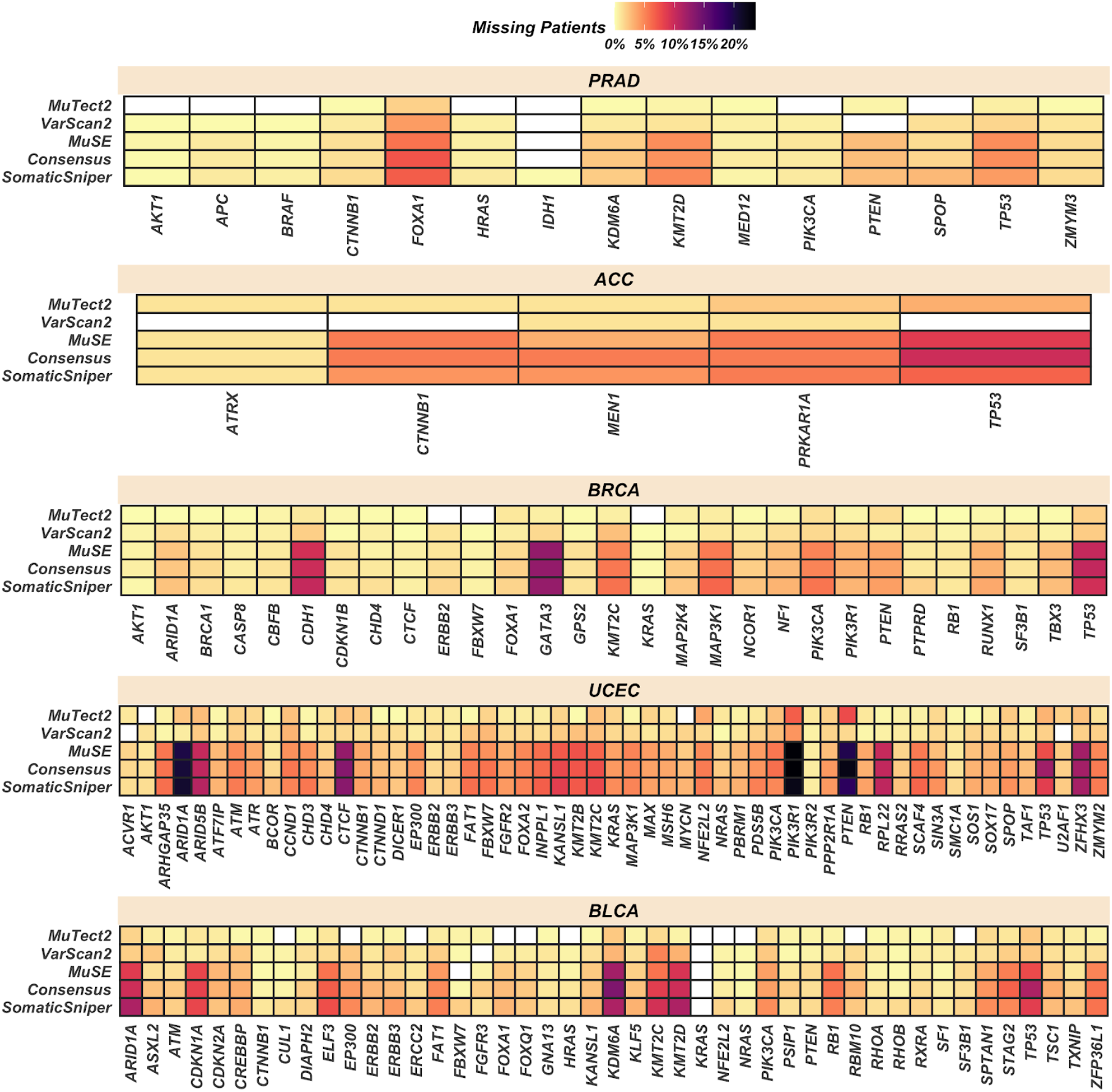
Detection of somatic mutations in cancer driver genes. Patients with cancer driver genes differing mutation status depending on the variant call set used. Union call set is used as reference to consider all possible samples bearing mutations in a given cancer driver gene. The percentage of these Union call set samples missed by each variant call set is shown. Only the PanCancerAtlas -MC3- project cancer driver genes are shown.

Overall, as expected, there is a correlation between the total number of mutations called by each method and the number of mutations identified in cancer driver genes. MuTect2 and specially VarsCan2 detected more mutations in cancer driver genes than SomaticSniper, MuSe and the Consensus. Interestingly, driver genes detected in multiple cancer types, such as TP53, ARID1A or PTEN, also showed larger differences across call sets. In particular, TP53 is the gene with the largest difference in the number of patients with detected somatic mutations, with 5-15% patients that vary their TP53 mutational status across cancers depending on the variant call set selected. Other cancer driver genes with significant numbers of patients with differing somatic mutation status, between 10% and 15%, are ARID1A, KDM6A, KMT2C and KMT2D in BLCA or CDH1 and GATA3 in BRCA. In terms of cancer types, UCEC shows the largest variation across call sets with more than 20% of patients with diverging mutation status in ARID1A, PIK3R1 and PTEN depending on which mutation call set is used. Importantly, none of the somatic variant call sets had all the mutations in all cancer driver genes, suggesting that we need to use multiple variant callers to ensure that we are detecting all cancer driver mutations.

### 3.3 Mutational signatures

The analysis of mutational signatures is important to understand the biological mechanisms underlying somatic mutations such as defective DNA repair, mutagenic exposures, DNA replication infidelity or enzymatic DNA modifications. These mutational processes have implications in the understanding of cancer aetiology and may inform patient treatment.

We analyzed the mutational signatures in our cohorts focusing on those signatures that have been proved to contribute mutations to the corresponding cancer types^40^ **(Figure S7)**.

Overall, we detected all the expected mutational signatures in all cancer types regardless of the variant calling tool or strategy used. As expected, the mutational signatures contributing the most mutations to individual tumor genomes were SBS1, SBS2, SBS5, SBS13 and SBS40 (**Figure 5**).

**Fig. 5.**
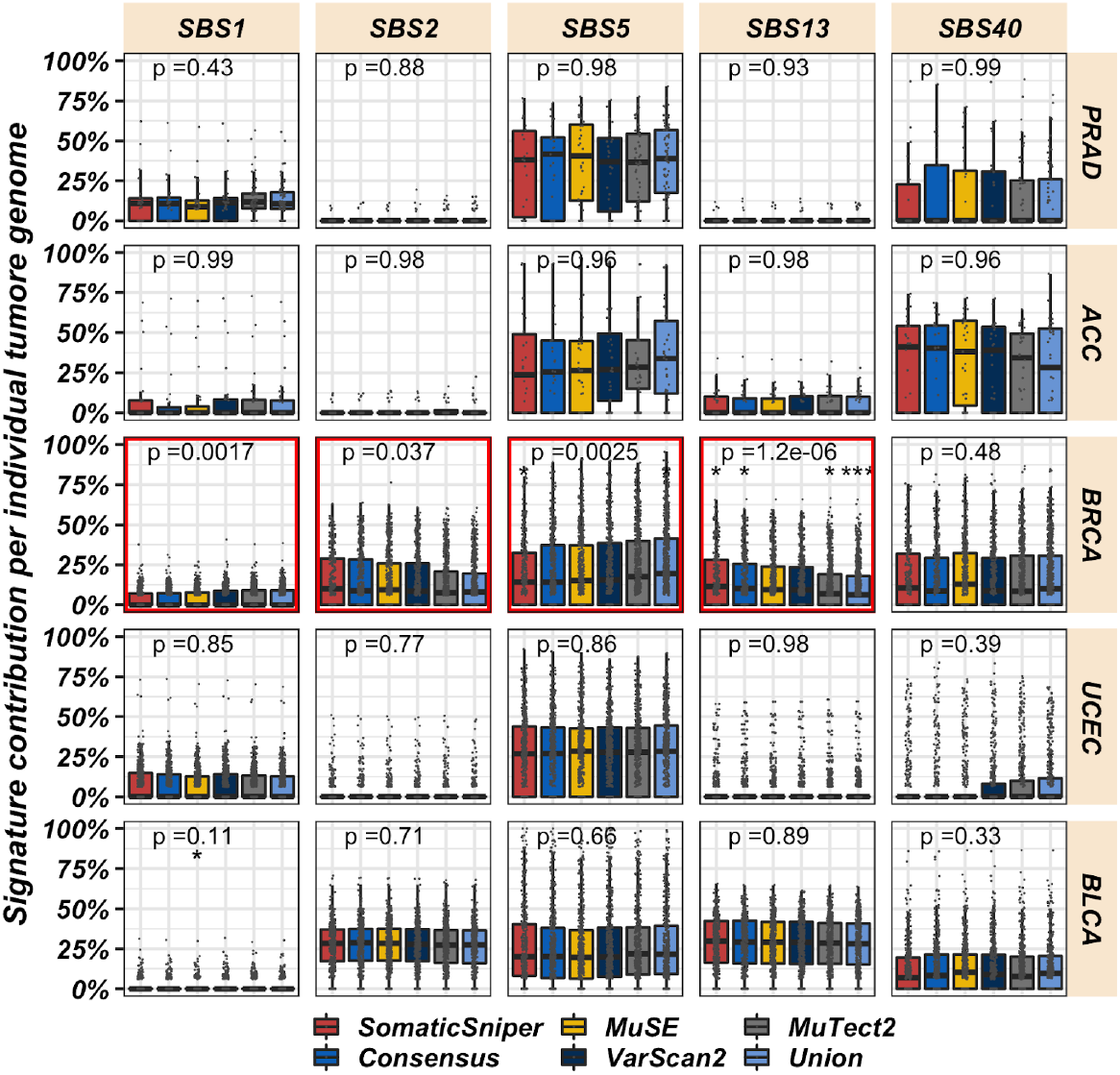
The percentage of mutations contributed by each mutational signature to individual tumor genomes. The mutational signatures contributing the most mutations to individual tumor genomes are shown. Those signatures showing significant differences in their contribution to individual tumor genomes depending on the variant call set used are squared in red. Global p.value was computed with Kruskal-Wallis test comparing all variant calling strategies. Pairwise comparisons were performed with Wilcoxon test comparing each strategy (reference) against all others (* = p.value < 0.05, ** = p.value < 0.01, *** p.value < 0.001). See also Figure S7.

We observe SBS5 and SBS40 as flat signatures contributing to multiple types of cancer, although their proposed aetiology remains unknown. Furthermore, SBS5, SBS40 and SBS1 mutations have been proved to correlate with age. Specifically, SBS1 may reflect the number of cell divisions a cell has undergone. On the other hand, cancers with high APOBEC activity, specially BLCA and to a lesser extent BRCA, show an increase in the mutational burden of SBS2 and SBS13, both of them related to the APOBEC family of cytidine deaminases activity.

For the majority of signatures and cancer types no differences are found among variant call sets. We only found significant differences in BRCA. In this cancer type, SBS13 contribution shows the largest difference between variant call sets (Kruskal-Wallis p value = 1.2e-06), followed by SBS1 (p < 0.002), SBS2 (p < 0.05) and SBS5 (p < 0.003). We found no statistically significant differences in the quantification of mutational signatures depending on the variant call set used in any other cancer cohort. We would like to emphasize that one of the main sources of false positives callings are germline mutations in CpG sites that are miscalled as somatic. Hence, the lack of significant differences in SBS1 (characterized by C>T mutations at NCG trinucleotides; N being any base) results is relevant. Overall, it seems that mutational signatures are pretty robust to variant calling decisions, with the exception of BRCA.

### 3.4 Differences in clinically actionable mutations depending on the variant calling strategy

Another important goal of the analysis of somatic cancer genomes is the identification of clinically actionable variants. These are somatic variants that help oncologists and physicians decide whether they should give a treatment to a cancer patient, as they are associated with sensitivity or resistance. Therefore, properly assessing the presence of such variants in the genome of cancer cells is of ultimate clinical importance. Thus, we used the Cancer Genome Tnterpreter^41^ to detect and classify variants as biomarkers of drug response according to different levels of clinical evidence.

We found many differences in the number of clinically actionable variants depending on which variant caller is used (**Figure 6**). As expected from their total number of mutations, the mutation sets from MuTect2, Union and VarScan2 have between 5-15% more patients with at least one clinically actionable mutation compared to Consensus, MuSE and SomaticSniper call sets. These results reflect the fact that the larger the variant call set is, the more likely it is to find a clinically actionable target.

**Fig. 6.**
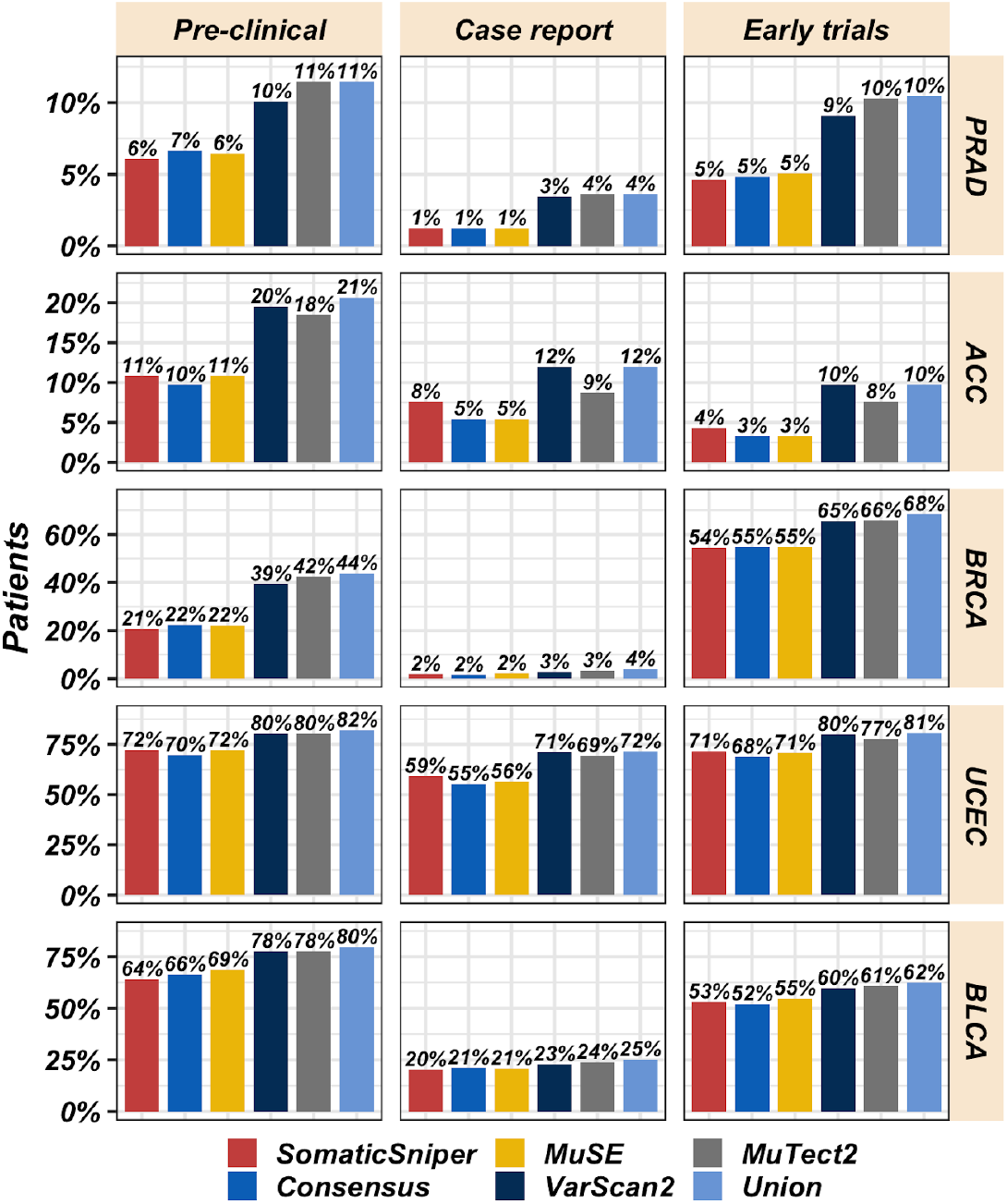
Clinically actionable mutations. Percentage of patients with at least one clinically actionable mutation identified by Cancer Genome Interpreter (CGI) with a given variant call set at various stages of approval. See also Figure S8.

We found larger differences among variant call sets in BLCA, BRCA and UCEC. Additionally, we found the most important differences in BRCA with MuTect2, Union and VarScan2 call sets identifying 10% and 20% more patients bearing at least one clinically actionable mutation within the early trials and preclinical evidence levels respectively.

The actionable variants that contribute to the most number of samples are TP53 mutations, found in 30% and 50% of BRCA and BLCA patients respectively (**Figure S8**). PTK3CA mutations affect more than 30% and 40% of BRCA and UCEC patients respectively. Moreover, we found larger differences in UCEC patients bearing PTEN clinically actionable mutations, with 40% to 60% of patients bearing this actionable mutation depending on the variant calling strategy. This 20% difference is important since these mutations may predict sensitivity to PT3K pathway inhibitors, mTOR and PARP inhibitors.

## 4 Discussion

The analysis of sequencing data from cancer genomes is critical, among others, to understand cancer aetiology, identify the events driving the transformation of healthy cells into cancerous ones or guide the treatment of cancer patients. Each of these analyses rely on the proper identification of true somatic variants in the cancer genome, which can be done with many different computational tools. However, to the best of our knowledge, we currently do not understand the impact of variant calling approaches to the results of secondary analyses of cancer somatic mutations.

Here we have quantified the impact of changing variant calling tools or strategies in three different secondary analyses across five different cancer types. We have shown that variant calling decisions lead to different outcomes in the identification of cancer driver genes, mutational signatures and clinically actionable variants.

Regarding cancer driver genes, while the performance of each variant calling tool or strategy can vary depending on the cancer type, the overall results suggest that one will get the best results using the mutations from either MuTect2 alone or the Union of all variant calling tools. These strategies lead to the detection of more driver genes than the rest of variant calling strategies. The result of the Union variant call set was a surprise for us. We initially expected that the likely high number of false positive somatic mutations in the Union call set would lower the predictive power of the cancer driver detection tools in intOgen, but this was not the case. In fact, the most common approach to combine somatic variant call sets is the Consensus^5^, but this led to worse overall results to detect cancer driver genes. This shows how starting from a call set with supposedly lots of false positive calls, one can still correctly identify the majority of cancer driver genes in a cohort.

Importantly, we have also found differences in the detection of somatic mutations in cancer driver genes. In some cases, a specific cancer driver gene mutation status could differ in more than 20% of patients depending on the variant call set used. This result suggests that it is important to use, at least, more than one variant calling tool to analyze cancer genomes. Otherwise a significant number of mutations in cancer driver genes can be missed.

Regarding mutational signatures, we have shown that their analysis is mostly robust to the variant call set selected. However, we observed significant differences in the signatures contributions to individual tumor genomes for four of the most common signatures in breast cancer: SBS1, SBS2, SBS5, SBS13.

If the goal of the analysis of the somatic genome is to find clinically actionable mutations, we need to be aware that there are considerable differences depending on the somatic mutation calling used. On average, 10% of patients with at least one clinically actionable target can either be detected or not depending on the variant calling strategy.

We acknowledge several limitations in our study. For example, we are not considering results for copy number and structural variants in the mutation call sets. Additionally, the limited availability of experimentally validated variants with orthogonal sequencing prevents us from extracting final conclusions regarding which variant caller performance is more robust.

We hope this study will help researchers understand how variant calling decisions might impact their results. It is important to account for the clinical implications that variant calling decisions have on different downstream analyses, especially in such important aspects of cancer genomics like driver genes, mutational signatures and the identification of actionable variants. Moreover, we aim this study will help guide variant calling design while considering the needs and goals of the different research projects. For example, as explained above, the most common strategy to combine the output of multiple variant calling tools, the Consensus, seems to have some shortcomings, even when compared to the simpler Union strategy.

Finally, while the optimal variant calling strategy seems to depend on both, the secondary analysis of interest as well as the cancer type being studied, the single recommendation that we believe can be applied in all circumstances is to use, at least, more than one variant calling tool and test the results of any secondary analysis in the different variant call sets. This would give researchers a sense of how much their results might vary depending on the variant calling and whether additional efforts into running other variant calling tools are necessary or not.

## Acknowledgements

We would like to thank the patients that donated the samples for The Cancer Genome Atlas, without them this work would not be possible. We would also like to thank Abel Gonzalez-Perez for his valuable discussions and insights.

*Conflict of Interest:* none declared.

