## Supplemental Figures for "The consequences of variant calling decisions in secondary analyses of cancer genomics data"

**Figure S1. TCGA projects characteristics.** (A) Biological sex distribution. Coloured bars represent the five cohorts we have analysed. (B) Purity distribution. (C) Mutational load distribution. Gray box plots represent the five cohorts we have analysed.

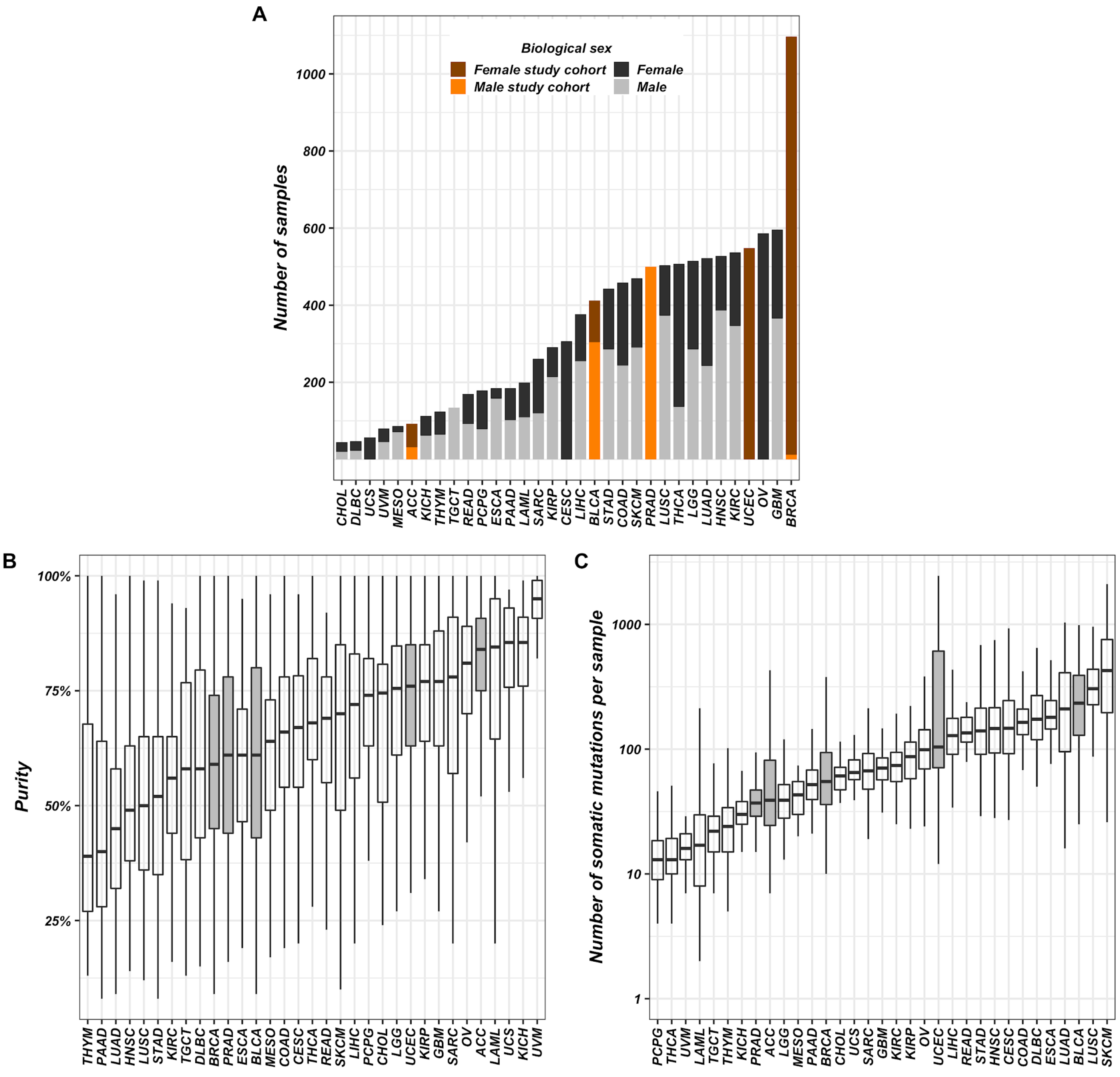

**Figure S2. Intersection of mutation calls across variant calling strategies by cancer type.** This UpSetR plot shows the number of variants uniquely called by one tool (single point) and variants called by different tools (linked points). Top bar-plot indicates intersection size. Bottom left plot indicates call set size. (A) PRAD. (B) ACC. (C) BRCA. (D) UCEC. (E) BLCA.

A **Prostate adenocarcinoma (PRAD)**

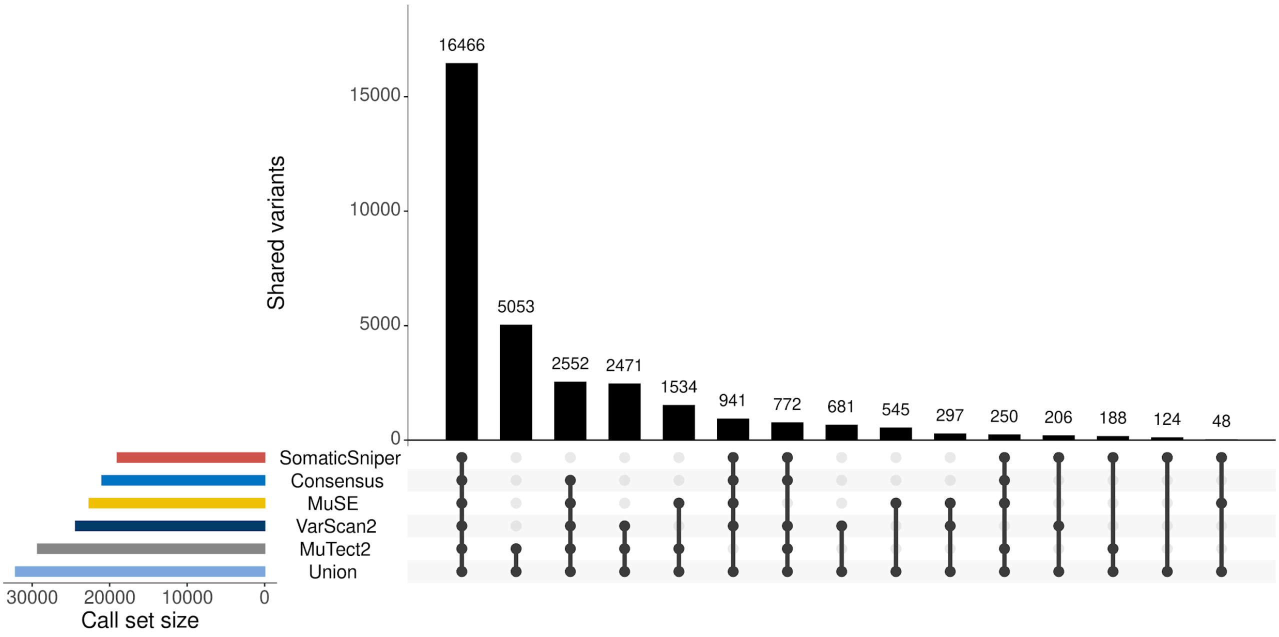


B **Adrenocortical carcinoma (ACC)**

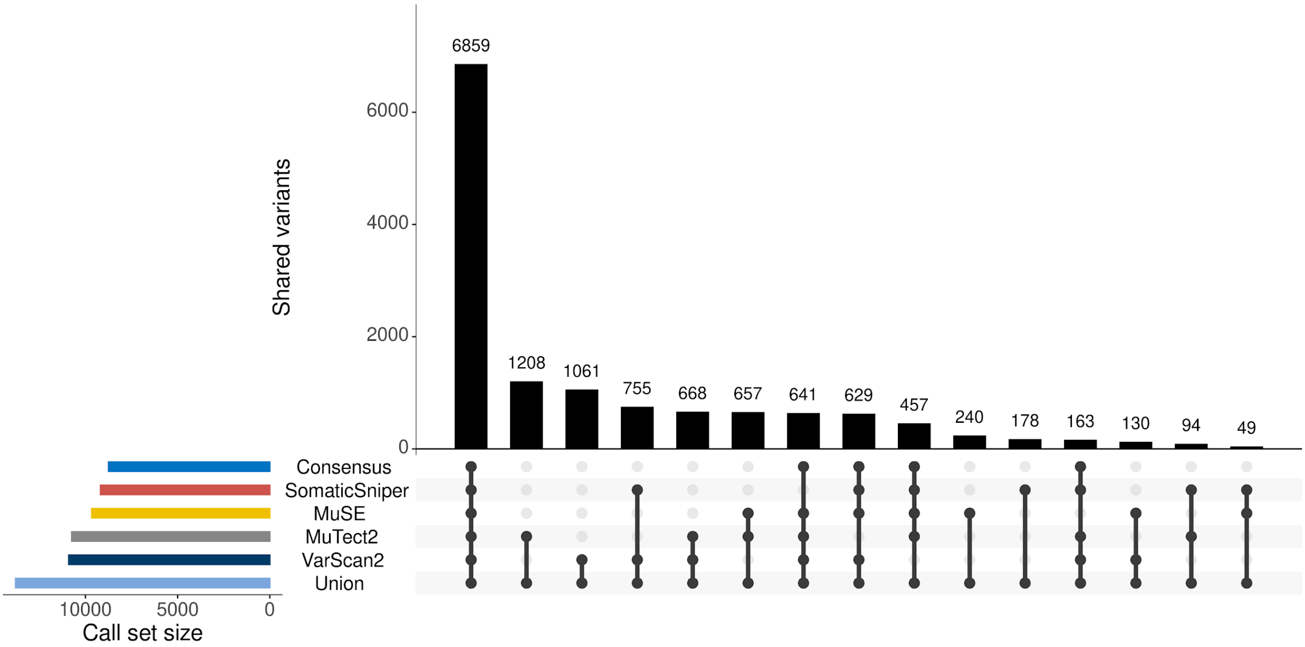

C Breast invasive carcinoma (BRCA)

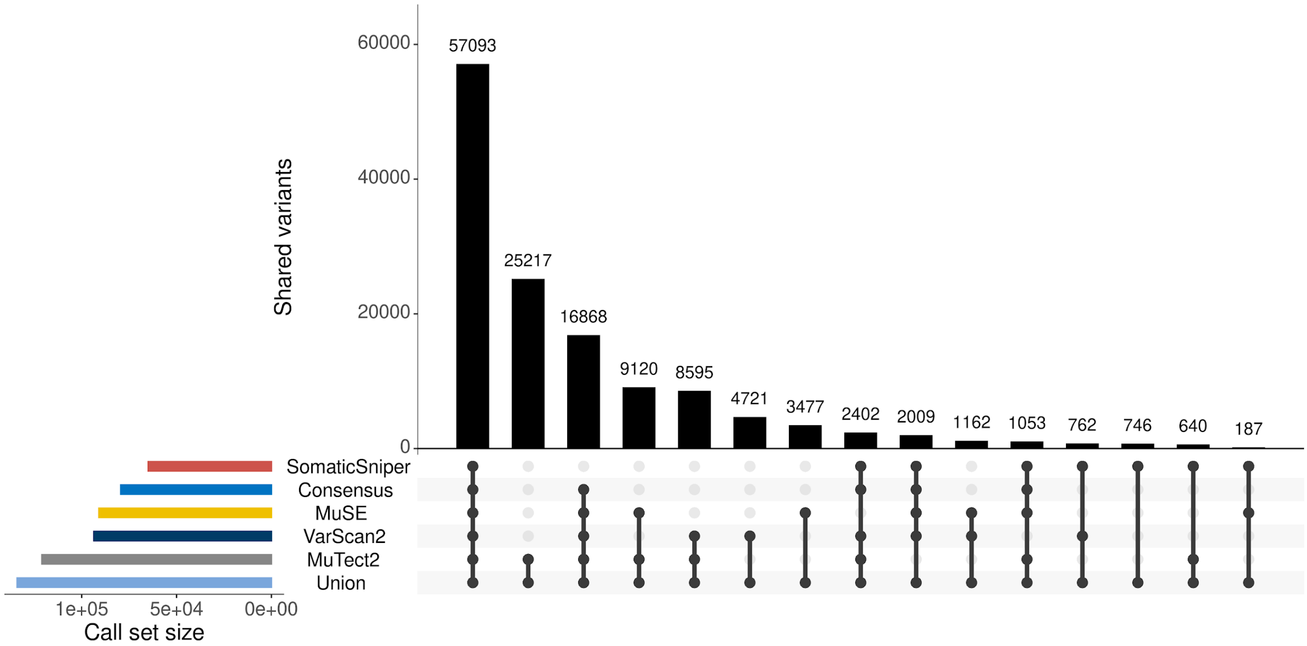


D Uterine corpus endometrial carcinoma (UCEC)

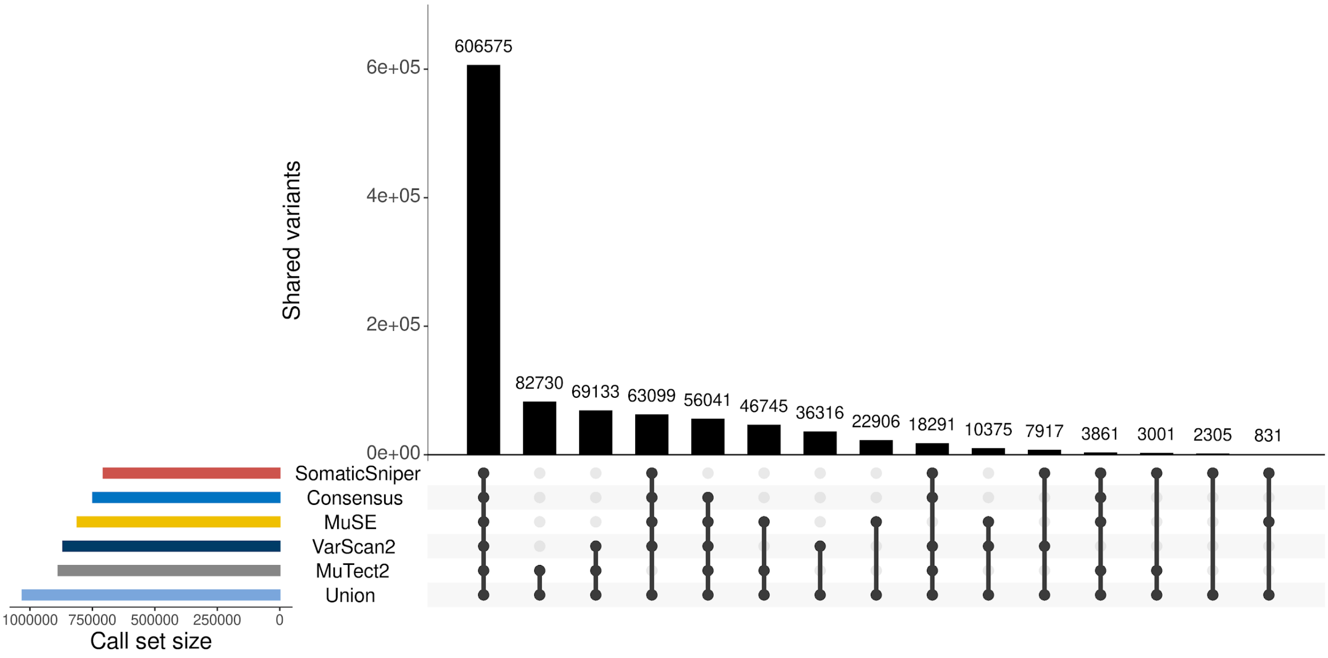

E **Bladder urothelial carcinoma (BLCA)**

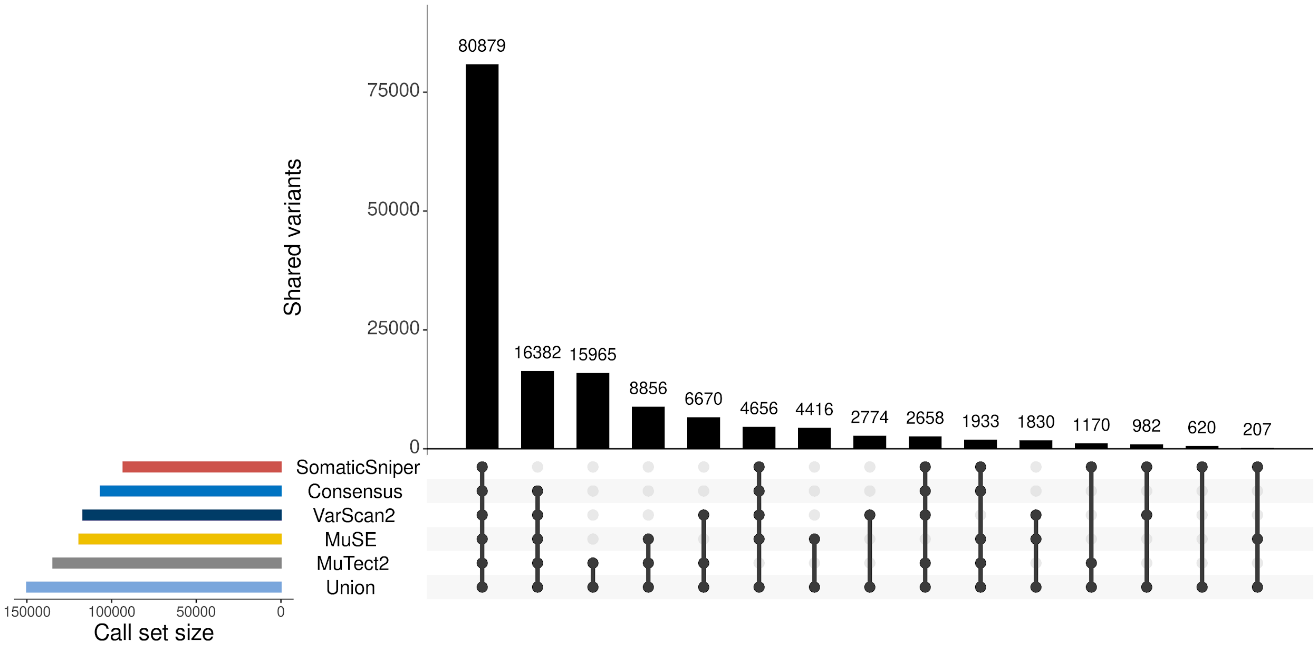

**Figure S3. Correlation between tumor sample purity and Union call set variants identified.** This scatter plot shows the spearman correlation between tumor sample purity and the percent of all possible somatic variants (Union call set) called by each variant calling strategy. All five cancer types results are shown together. Dots represent tumor samples. Shaded area indicates 95% bootstrapped confidence interval.

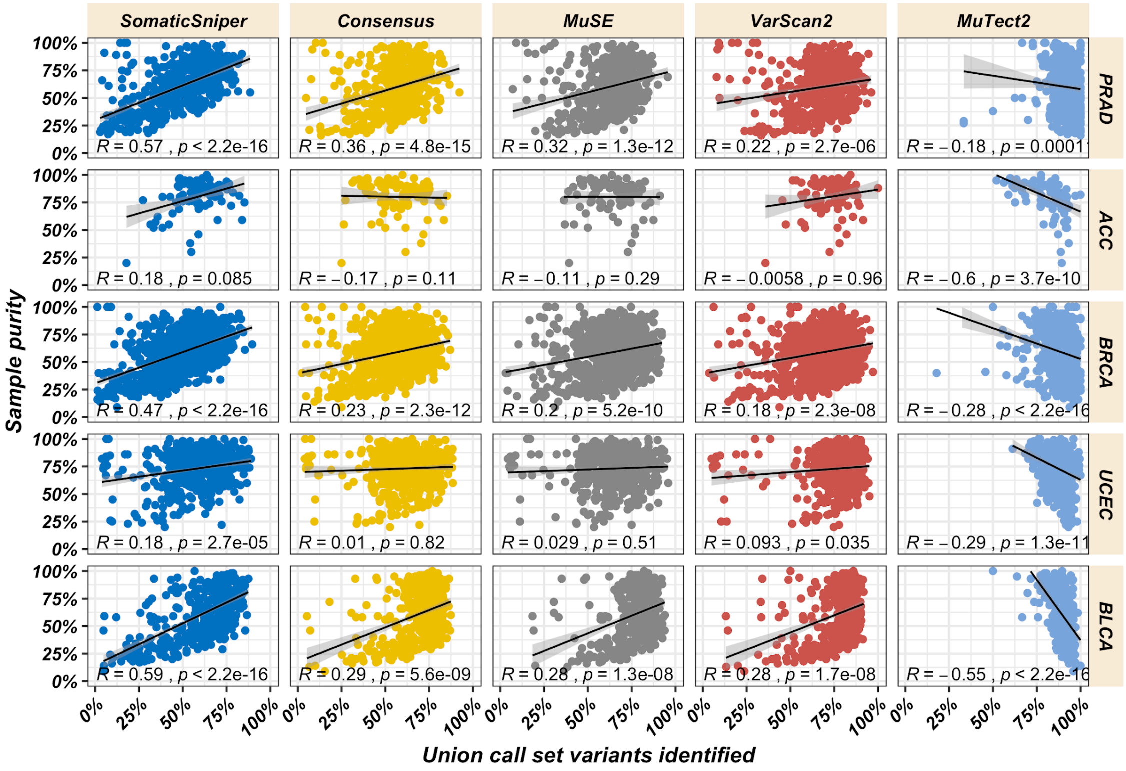

**Figure S4. Correlation between tumor sample purity and tumor stage**. This box plot shows the correlation between tumor sample purity and tumor sample stage[44]. AJCC (American Joint Committee on Cancer) stages for ACC, BLCA and BRCA. Clinical stage for UCEC. No stage information was available for PRAD. Dots represent tumor samples.

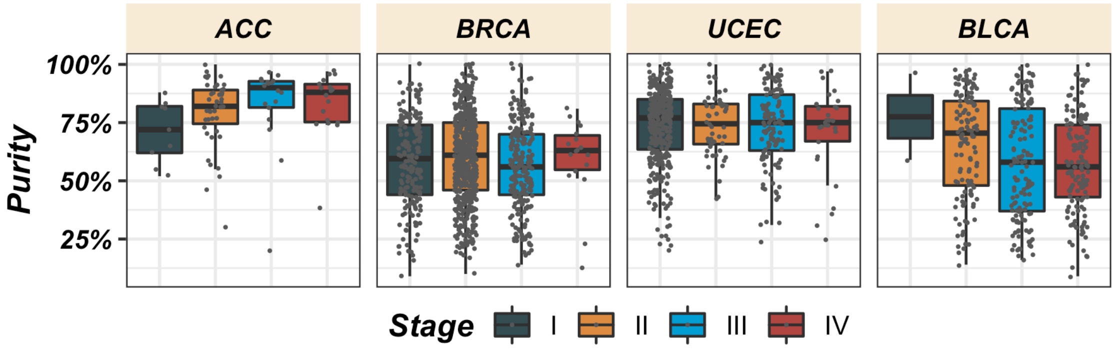

**Figure S5. Variant allele frequency (VAF) distribution.** Comparison of the VAF range spanned by each call set. The VAF was adjusted by purity and ploidy[43]. Global p.value was computed with Kruskal-Wallis test comparing all variant calling strategies. Pairwise comparisons were performed with Wilcoxon test comparing each strategy (reference) against all others (* = p.value < 0.05, ** = p.value < 0.01, *** = p.value < 0.001, **** = p.value < 0.0001).

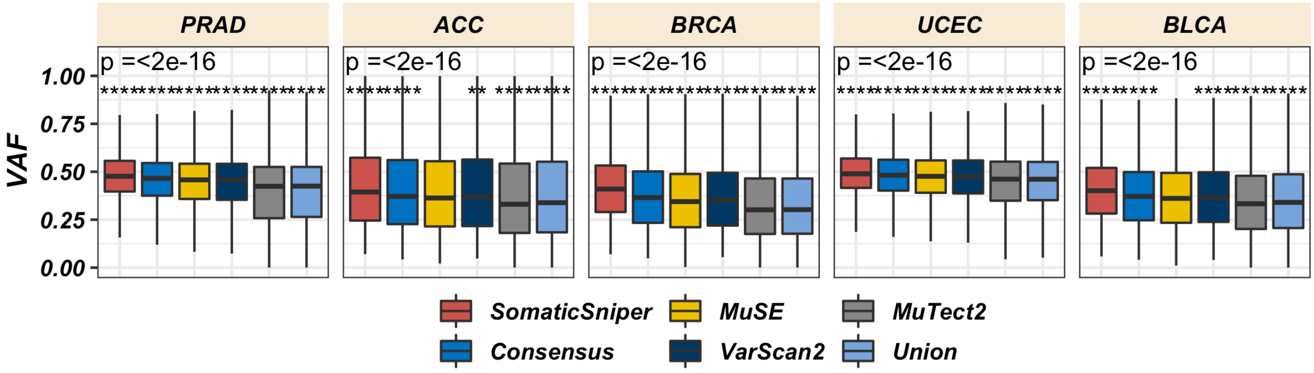

**Figure S6. Performance metrics of the different variant calling strategies when detecting cancer driver genes with intOGen.** The results for each cancer type are shown. (A) performance metrics when benchmarking against intOGen truth set of known cancer driver genes. (B) performance metrics when benchmarking against PanCancerAtlas projects -MC3- truth set of known cancer driver genes. FDR: False Discovery Rate.

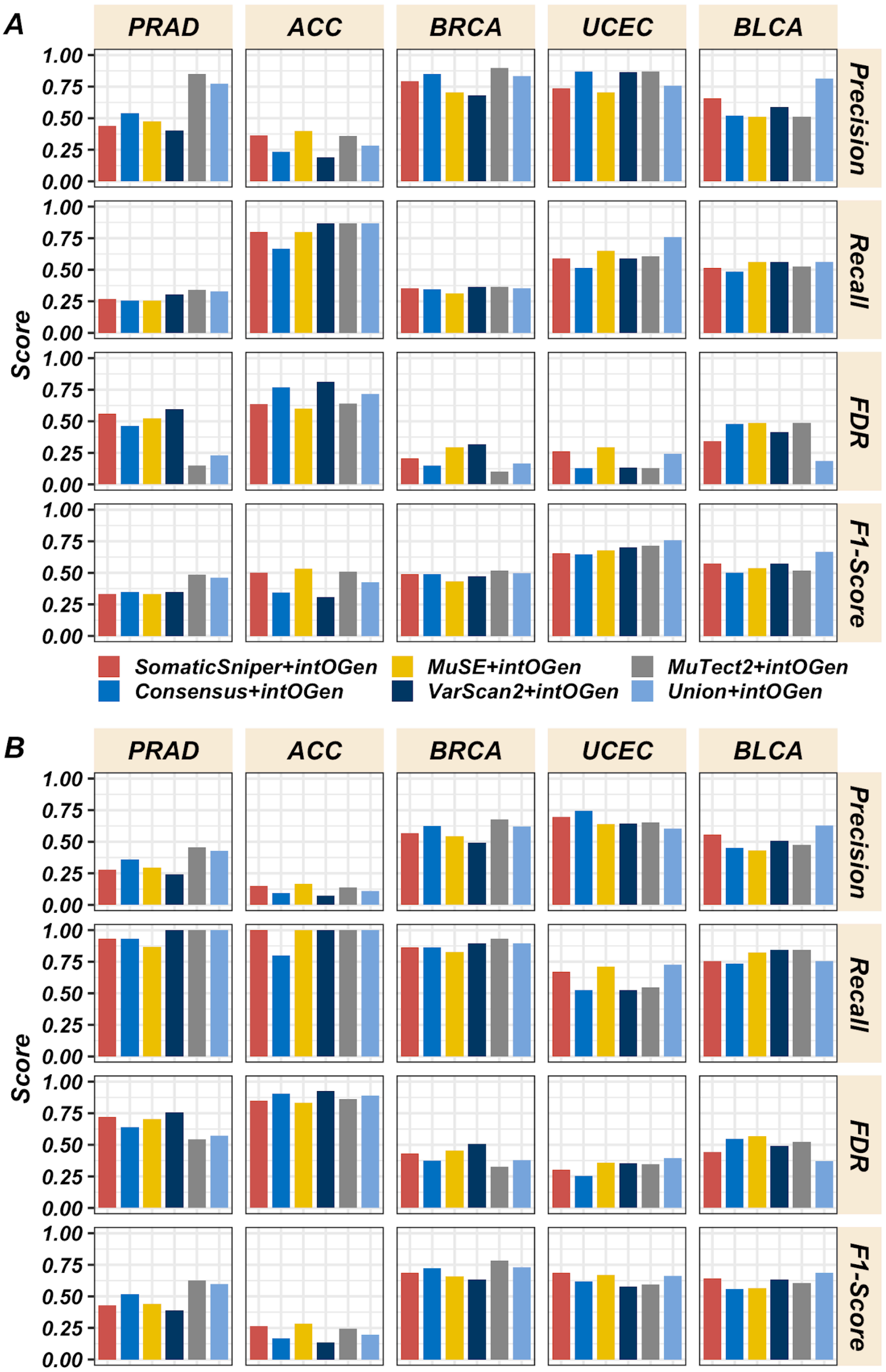

**Figure S7. The percentage of mutations contributed by each mutational signature to individual tumor genomes.** Signatures that have been proved to contribute mutations to the corresponding cancer type[40] are shown. Those signatures showing significant differences in their contribution to individual tumor genomes depending on the variant call set used are squared in red. Global p.value was computed with Kruskal-Wallis test comparing all variant calling strategies. Pairwise comparisons were performed with Wilcoxon test comparing each strategy (reference) against all others (* = p.value < 0.05, ** = p.value < 0.01, *** = p.value < 0.001). (A) PRAD. (B) ACC. (C) BRCA. (D) UCEC. (E) BLCA.

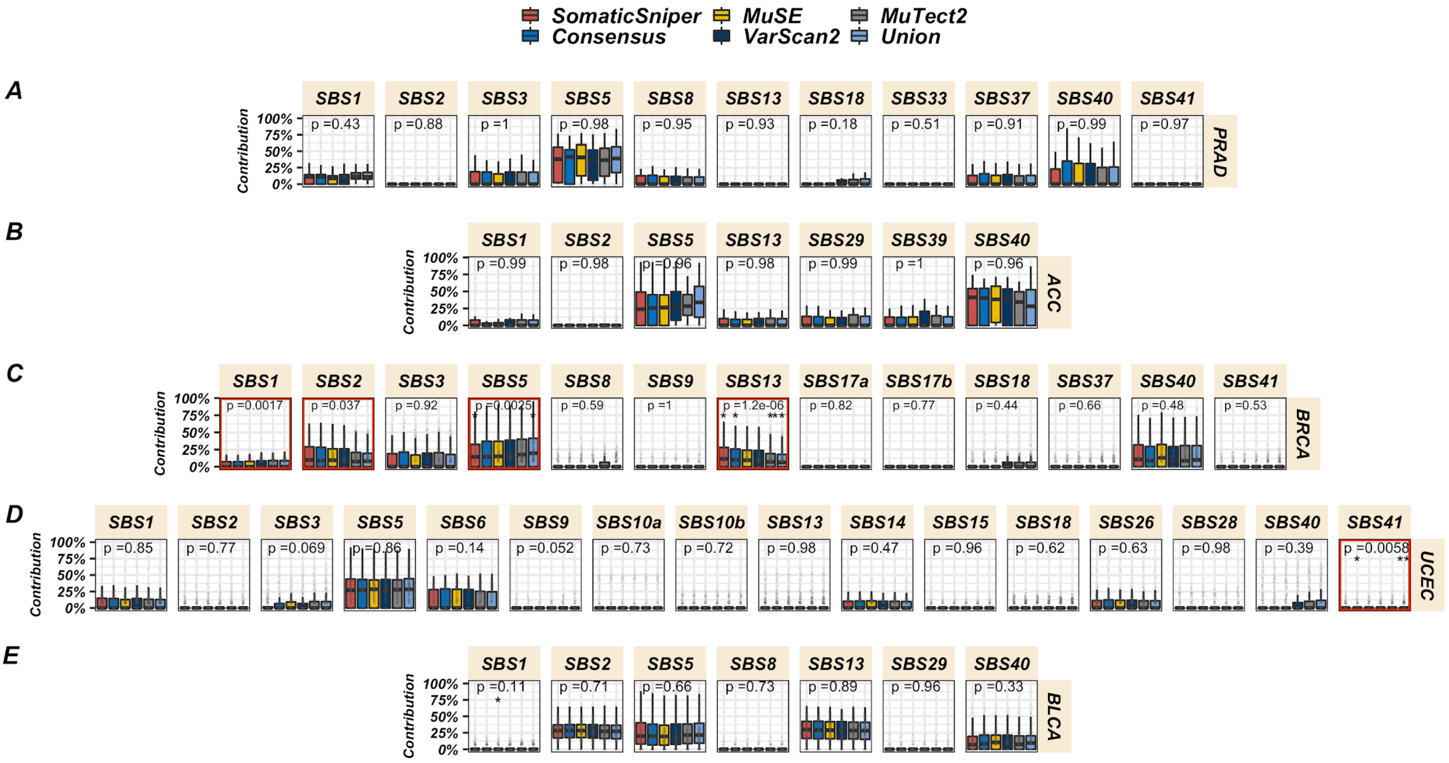

**Figure S8. Cancer driver genes bearing clinically actionable mutations.** Percentage of patients with at least one clinically actionable mutation, regardless of stage of approval, in a given cancer driver gene identified by Cancer Genome Interpreter (CGI) with a given variant call set. Most frequently mutated driver genes are shown. Only those cancers with larger differences among variant call sets are displayed.

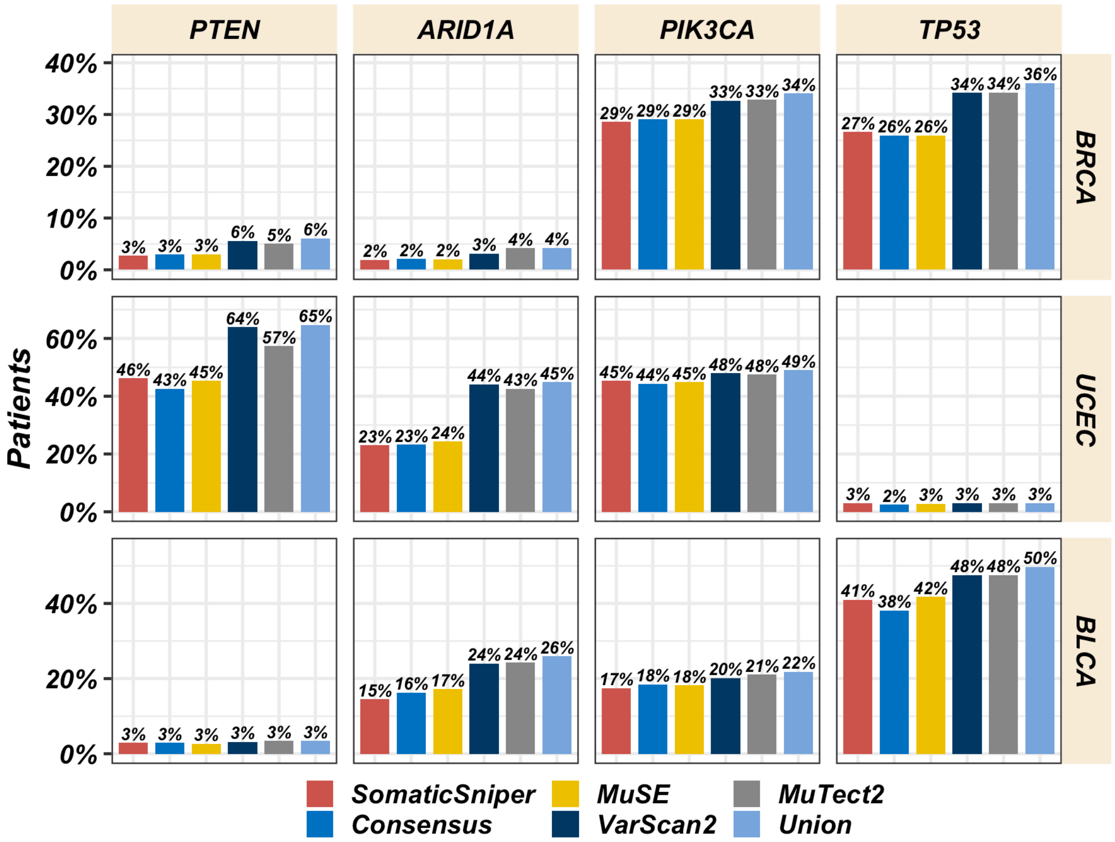

**Supplemental Table S1 (attached Excel XLSX file): intOGen results.** Cancer driver genes detected (represented by 1) and non-detected (represented by 0) by intOGen with a given variant call set in each cancer cohort. In the case of intOGen and MC3[5] truth sets, 1 stands for those cancer driver genes present in the truth set and 0 for those genes not present in the truth set.
